# The answer lies in the energy: how simple atomistic molecular dynamics simulations may hold the key to epitope prediction on the fully glycosylated SARS-CoV-2 spike protein

**DOI:** 10.1101/2020.07.22.214254

**Authors:** Stefano Serapian, Filippo Marchetti, Alice Triveri, Giulia Morra, Massimiliano Meli, Elisabetta Moroni, Giuseppe A. Sautto, Andrea Rasola, Giorgio Colombo

## Abstract

Betacoronavirus SARS-CoV-2 is posing a major threat to human health and its diffusion around the world is having dire socioeconomical consequences. Thanks to the scientific community’s unprecedented efforts, the atomic structure of several viral proteins has been promptly resolved. As the crucial mediator of host cell infection, the heavily glycosylated trimeric viral Spike protein (S) has been attracting the most attention and is at the center of efforts to develop antivirals, vaccines, and diagnostic solutions.

Herein, we use an energy-decomposition approach to identify antigenic domains and antibody binding sites on the fully glycosylated S protein. Crucially, all that is required by our method are unbiased atomistic molecular dynamics simulations; no prior knowledge of binding properties or ad hoc combinations of parameters/measures extracted from simulations is needed. Our method simply exploits the analysis of energy interactions between all intra-protomer aminoacid and monosaccharide residue pairs, and cross-compares them with structural information (i.e., residueresidue proximity), identifying potential immunogenic regions as those groups of spatially contiguous residues with poor energetic coupling to the rest of the protein.

Our results are validated by several experimentally confirmed structures of the S protein in complex with anti- or nanobodies. We identify poorly coupled sub-domains: on the one hand this indicates their role in hosting (several) epitopes, and on the other hand indicates their involvement in large functional conformational transitions. Finally, we detect two distinct behaviors of the glycan shield: glycans with stronger energetic coupling are structurally relevant and protect underlying peptidic epitopes; those with weaker coupling could themselves be poised for antibody recognition. Predicted Immunoreactive regions can be used to develop optimized antigens (recombinant subdomains, synthetic (glyco)peptidomimetics) for therapeutic applications.

## Introduction

The novel coronavirus SARS-CoV-2, the etiological agent of COVID-19 respiratory disease, has infected millions of people worldwide, causing more than 500000 deaths (as of June 30^th^, 2020) and extensive social and economic disruption. Given the pandemic status of the outbreak, social distancing measures cannot be sufficient any longer in containing it on a worldwide scale. This emergency calls for the development of strategies to rapidly identify pharmacological agents or vaccines as the only way to contain and combat the disease, restoring normal social conditions. Indeed, a number of currently ongoing trials focus on developing vaccines (see for instance https://www.nytimes.com/interactive/2020/science/coronavirus-vaccine-tracker.html) or on repurposing drugs already developed for other disorders ^1–4^.

SARS-CoV-2 is extraordinarily effective in exploiting the host’s protein machinery for replication and spreading. This is a characteristic that it shares with other members of the *Coronaviridae* family, all of which are characterized by a highly selective tropism that determines the onset of a variety of diseases in domestic and wild animals as well as in humans, including central nervous system affections, hepatitis and respiratory syndromes ^5–6^. As was the case with its human predecessors SARS-CoV and MERS, critical players regulating cell entry of SARS-CoV-2 are the homotrimeric viral Spike protein (S) (Figure 1) and the host cell docking point, the protein receptor angiotensin-converting enzyme 2 (ACE2)^7^. The CoV S protein is then cleaved by a series of serine proteases, including trypsin, cathepsins, elastase, the host type 2 transmembrane serine protease (TMPRSS2, DPP4 for MERS), and plasmin, which promote virus entry into epithelial cells ^4^.

**Figure 1.**
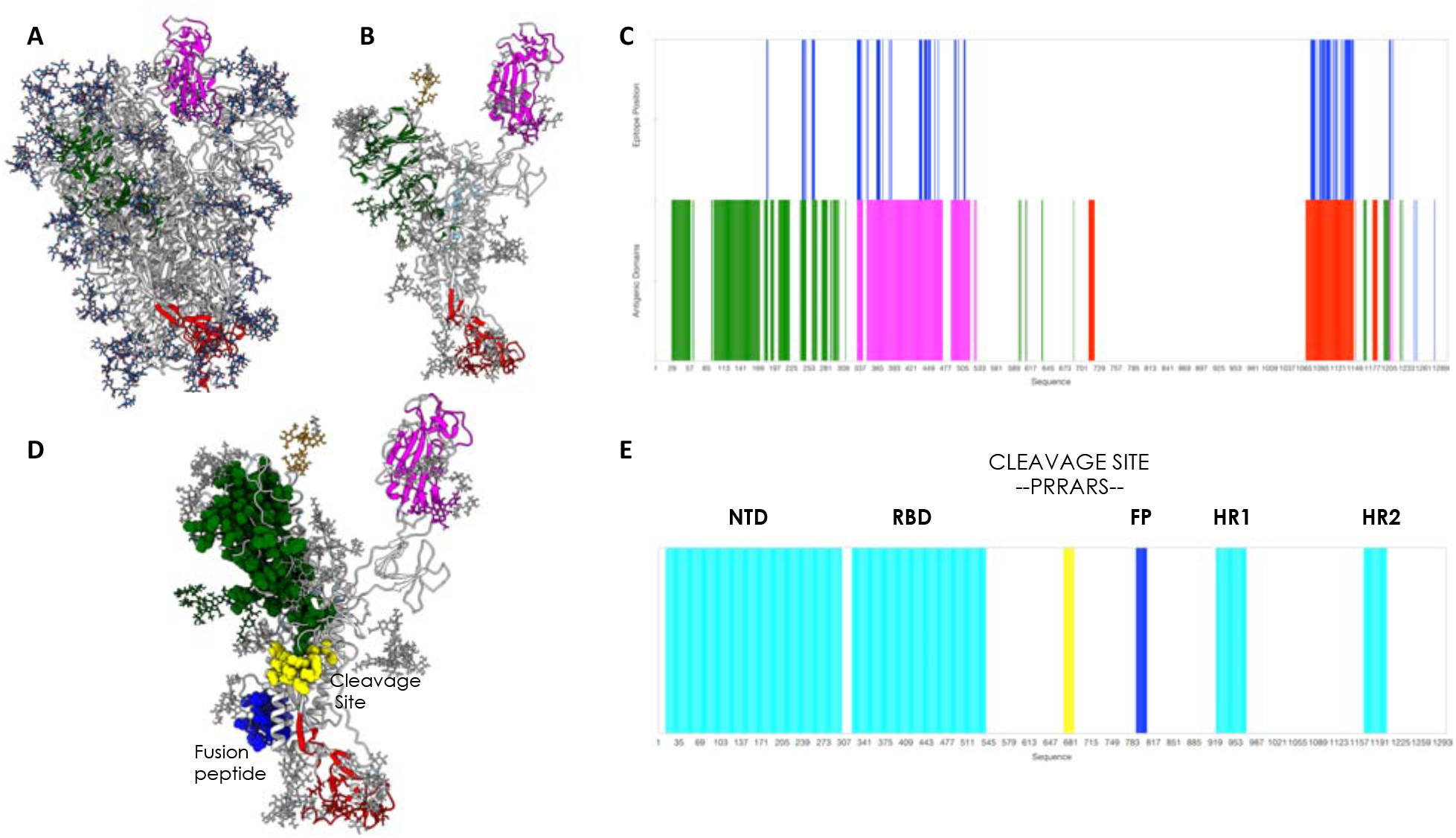
3D structure, glycosylation and location of antigenic domains and epitopes on SARS-CoV-2 fully glycosylated Spike protein. **A)** The starting, fully glycosylated Spike protein trimer. The coating oligosaccharides are colored in dark blue. The predicted antigenic domains are colored on the structure of one protomer. **B)** Isolated protomer with the most antigenic domains, detected via MLCE with the 15% cutoff, highlighted in colors: the antigenic part in the N-Domain is dark green; the part in the RBD is magenta; the part in the C-terminal domain is dark red. Oligosaccharides that define or are part of antigenic domains are also colored. Oligosaccharides that have a structural role and show strong energetic coupling to the protein are in white. **C)** The predicted antigenic sequences projected on the sequence of the protein. The bottom line reports the sequences defined as antigenic domains, with the same color code as in **B**. The top bar reports the location of peptidic epitopes identified with the most restrictive definition. **D)** Physical interaction between the boundaries of the predicted antigenic domain in the N-terminal region and the cleavage site of S. This subfigure also shows the physical proximity of the predicted C-terminal uncoupled region with the fusion peptide. **E)** Domain organization of the spike protein projected on the sequence. Numbering and domain definitions obtained from UNIPROT (https://www.uniprot.org/uniprot/P0DTC2)

Recent cryo-electron microscopy (cryo-EM) analyses allowed to precisely determine the structure of the full-length Spike protein in its trimeric form ^8–10^, and the structural basis for the recognition of the Spike protein’s Receptor Binding Domain (RBD; Figure 1) by the extracellular peptidase domain of ACE2 ^11^. In parallel, computational studies have started to provide atomically detailed insights into S protein dynamics and on the elaborate role of the diverse polysaccharide chains that decorate its surface, effectively shielding a large portion of it from the host ^12–14^.

This detailed dynamic and structural knowledge can set the stage for understanding the molecular bases of S protein recognition by the host’s immune system, providing information on which physicochemical determinants are required to elicit functional antibodies. Such understanding can then be exploited to design and engineer improved antigens based on S, for instance by identifying antigenic domains that can be expressed in isolation or short sequences (epitopes) that can be mimicked by synthetic peptides^15–20^: this would be a crucial first step in the selection and optimization of candidate vaccines and therapeutic antibodies, as well as in the development of serological diagnostic tools. Furthermore, knowledge acquired today about such recognition mechanisms could well mean that we are better prepared to tackle similar pandemics in the future by contrasting them more efficiently with the application of efficient and well-tested methods to new protein variants.

Here, we analyze representative 3D conformations of the full-length trimeric S protein in its fully glycosylated form (Figure 1), extracted from atomistic molecular dynamics (MD) simulations provided by the Woods group ^13^, in order to predict immunogenic regions.

To this end, a simple *ab initio* epitope prediction method that we previously developed for unmodified proteins ^21–23^ is optimized and extended to cover glycoproteins. The method is based on the idea that antibody-recognition sites (epitopes) may correspond to localized regions only exhibiting low-intensity energetic coupling with the rest of the structure; this lower degree of coupling would allow them to undergo conformational changes more easily, to be recognized by a binding partner, and to tolerate mutations at minimal energetic expense.

We show that our approach is indeed able to identify regions—also comprising carbohydrates—that recent structural immunology studies have shown to be effectively targeted by antibodies. On the same basis, our method predicts several additional potential immunogenic regions (currently still unexplored) that can then be used for generating optimized antigens, either in the form of recombinant isolated domains or as synthetic peptide epitopes. Finally, our results help shed light on the mechanistic bases of the large-conformational changes underpinning biologically relevant functions of the protein.

To the best of our knowledge, this approach is one of the first that permits to discover epitopes in the presence of glycosylation (an aspect that is often overlooked), starting only from the analysis of the physico-chemical properties of the isolated antigen in solution. Importantly, the approach does not require any prior knowledge of antibody binding sites of related antigenic homologs and does not need to be trained/tuned with data sets or ad hoc combinations of information on sequences, structures, SASA or geometric descriptors. The procedure is thus immediately and fully portable to other antigens.

## Results and Discussion

To reveal the regions of the S protein that could be involved in antibody (Ab) binding, we employ a combination of the Energy Decomposition (ED) and MLCE (Matrix of Low Coupling Energies) methods, which we previously introduced and validated^21–32^ and discuss in full in the Methods section.

Starting from 6 combined 400 ns replicas of atomistic molecular dynamics simulations of the fully glycosylated S protein in solution ^13^ (built from PDB ID: 6VSB ^8^), we isolate a representative frame from each of the three most populated clusters. ED and MLCE analyses of protein energetics assess the interactions that each aminoacid and glycan residue in S protomers establishes with every other single residue in the same protomer. In particular, we compute the nonbonded part of the potential energy (van der Waals, electrostatic interactions, solvent effects) implicitly, via an MM/GBSA calculation (molecular mechanics/generalized Born and surface area continuum solvation)^33^, obtaining, for a protomer composed of *N* residues (including monosaccharide residues on glycans), a symmetric *N* × *N* interaction matrix *M_ij_*. Eigenvalue decomposition of *M_ij_* highlights the regions of strongest and weakest coupling. The map of pairwise energy couplings can then be filtered with topological information (namely, the residue-residue contact map) to identify localized surface networks of low-intensity coupling (*i.e*., clusters of spatially close residue pairs whose energetic coupling to the rest of the structure is weak and not energetically optimized through evolution).

In our model, when these fragments are located on or near the surface, contiguous in space and weakly coupled to the protein’s ‘stability core’, they represent potential interaction regions (i.e., epitopes). Otherwise put, putative interacting patches are hypothesized to be characterized by nonoptimized intramolecular interactions. Actual binding to an external partner such as an Ab is expected to occur if favorable intermolecular interactions determine a lower free energy for the bound than the unbound state ^21, 23, 34^. Furthermore, minimal energetic coupling with the rest of the protein provides these subregions with greater conformational freedom to adapt to a binding partner and improves their ability to absorb mutations without affecting the protein’s native organization and stability in a way that could be detrimental for the pathogen: all these properties are indeed hallmarks of Ab-binding epitopes.

Once interacting vicinal residue pairs (*i, j*)are identified by cross-comparison with the residue-residue contact map (vide supra and Methods Section), identification of poorly coupled regions representing potential epitopes proceeds as follows. Residue pairs are firstly ranked in order of increasing interaction intensity (from weakest to strongest). Two distinct sets of energetically decoupled regions are then mapped by applying two distinct cutoffs (‘softness thresholds’) to the residue pair list: either from the first 15% or from the first 5% of the ranked pairs (i.e., the 5% or 15% of the residue pairs with the weakest energetic coupling).

The less restrictive 15% cutoff subdivides the full-length, fully folded S protein into potentially immunoreactive domains (see Figure 1B,C and Methods)^22, 25, 34^. The goal is to uncover regions that may normally be hidden from recognition by Abs in the native protein structure, but that can be experimentally expressed as isolated domains. Highly reactive neutralizing epitopes may in fact be present only in specific but transient conformations that are not immediately evident in the static X-ray and EM models of the protein or are not accessible even to large scale MD simulations. Presenting these (cryptic) regions for Ab binding through their isolated parent domains may prove more advantageous in developing new immunogens^22, 25^.

The more stringent epitope definition (5% cutoff) narrows the focus on those (smaller) intra-domain regions that could be directly involved in forming the interface with Abs, and that can then be used to guide the engineering of optimized antigens in the form of synthetic epitope peptidomimetics. In this context, to be defined as epitopes, the energetically uncoupled regions must be at least 6-residue long.

Upon using the larger cutoff value, a large cluster of energetically unoptimized residue pairs localize at the Receptor Binding Domain, correctly identifying it as the most antigenic unit in the S protein’s ‘RBD up’ protomer (Figure 1B,C magenta colored domain). Interestingly, when the lowest energy-coupled residue pairs are mapped onto the ‘up’ RBD of all three 3D structures isolated from MD, there is a large overlap with regions recognized by Abs and nanobodies (revealed by recent X-ray and cryo-EM structures). Importantly, for example, our calculation correctly identifies the binding region of mAb **CR3022**^35^, known to target a cryptic epitope that is exposed only upon significant structural rearrangement of the protein ^12^ (Figures 2 and Figure 4).

**Figure 2.**
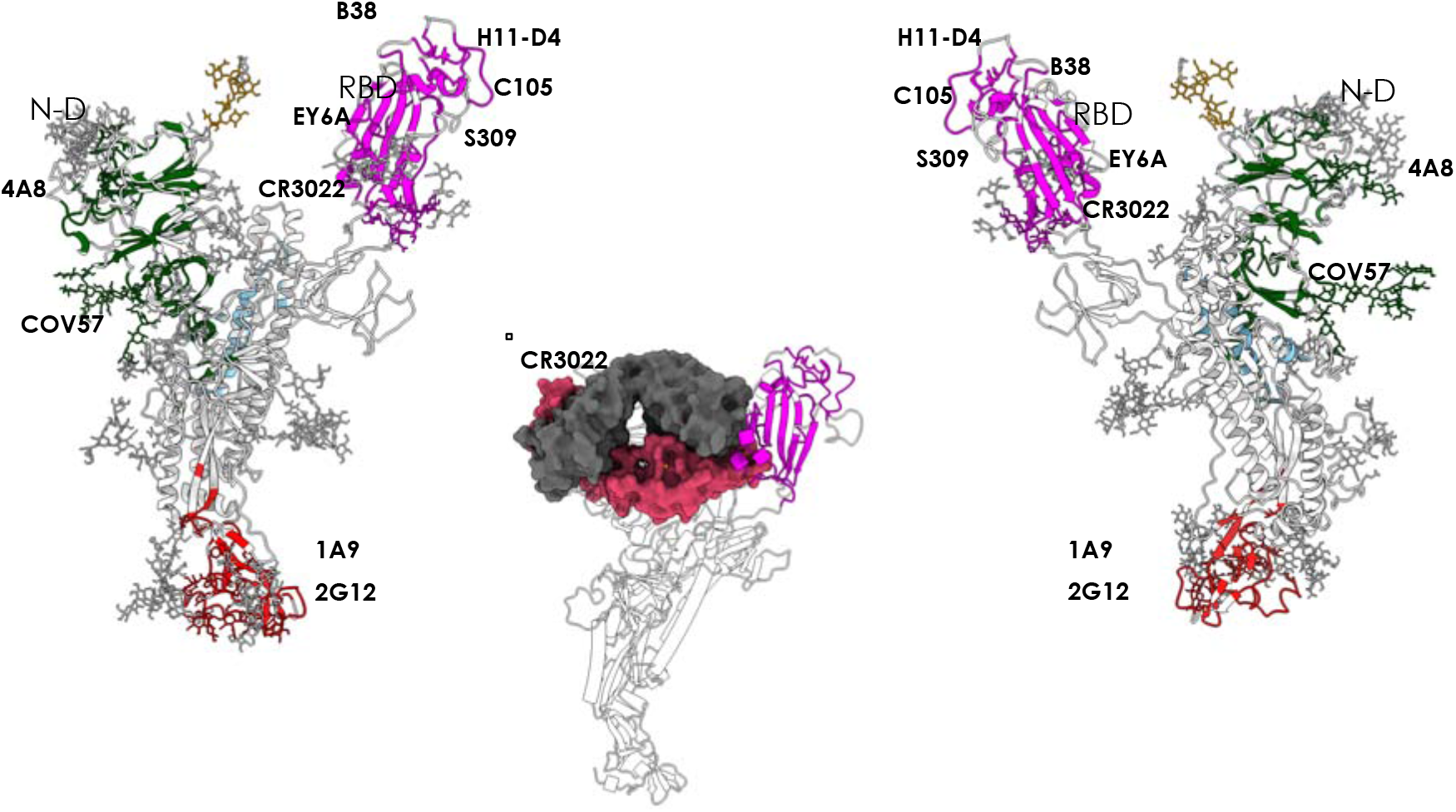
Antigenic domains and location of binding antibodies. The clusters of residues defining antigenic domains (dark green in the N-domain, magenta in the RBD, red in the C-terminal region) and the positions of the various antibodies whose structures and interactions in complexes with the full length protein have been described. The inset indicates the identification of the cryptic immunoreactive region binding CR3022.

**Figure 3.**
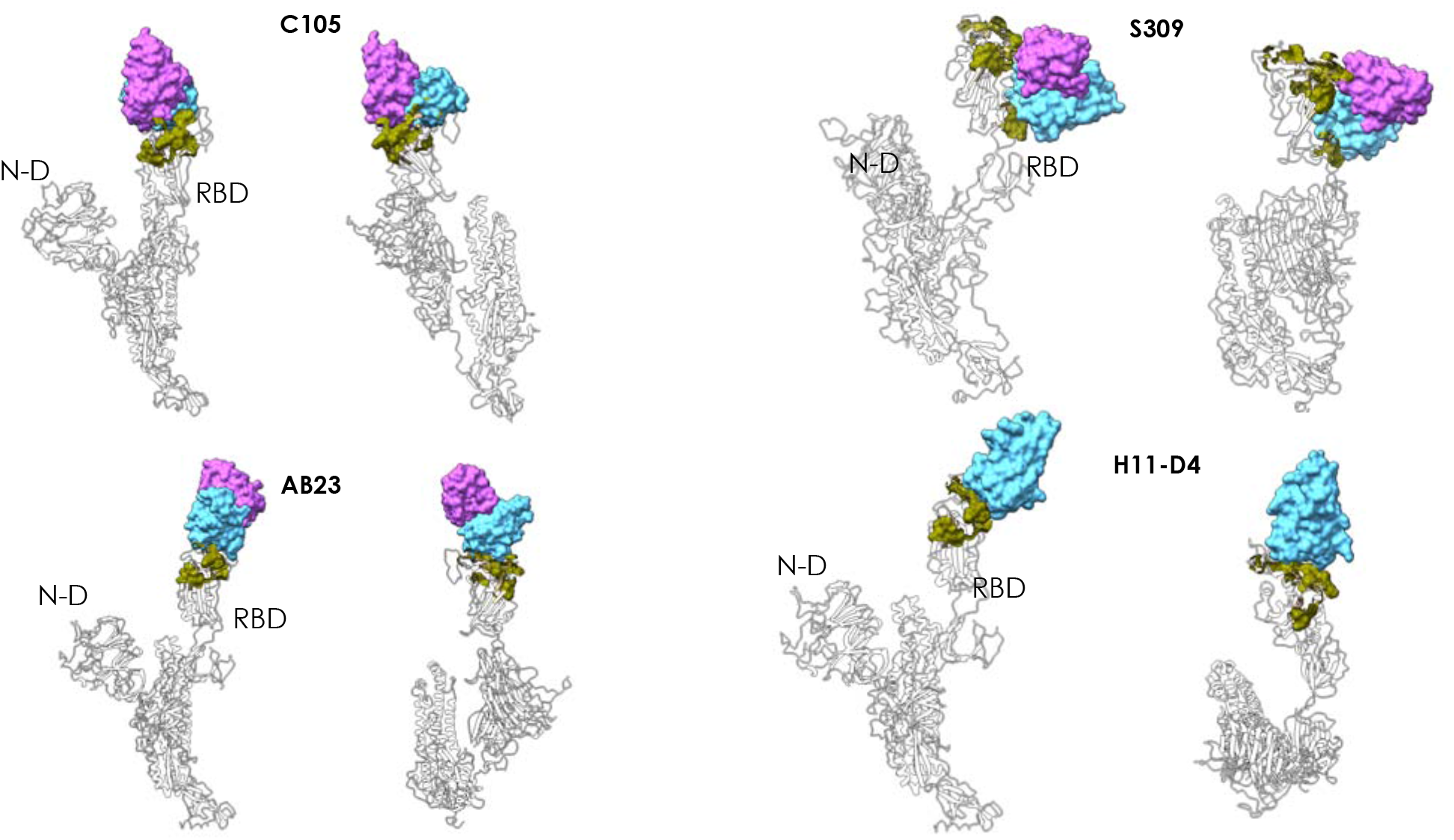
Peptidic epitopes predicted on the surface of the RBD using the restrictive definition of antigenic region and comparison with known Ab-complexes. The X-ray structures of the complexes between the various antibodies reported in the figure (C105, S309, AB23, and nanobody H11-D4) and the full length Spike protein are superimposed to the structure of the protomer used here for prediction. The green surfaces indicate the location of MLCE epitope predictions. The Fabs of the antibodies or of the nanobody are depicted as accessible surfaces in shades of blue.

**Figure 4.**
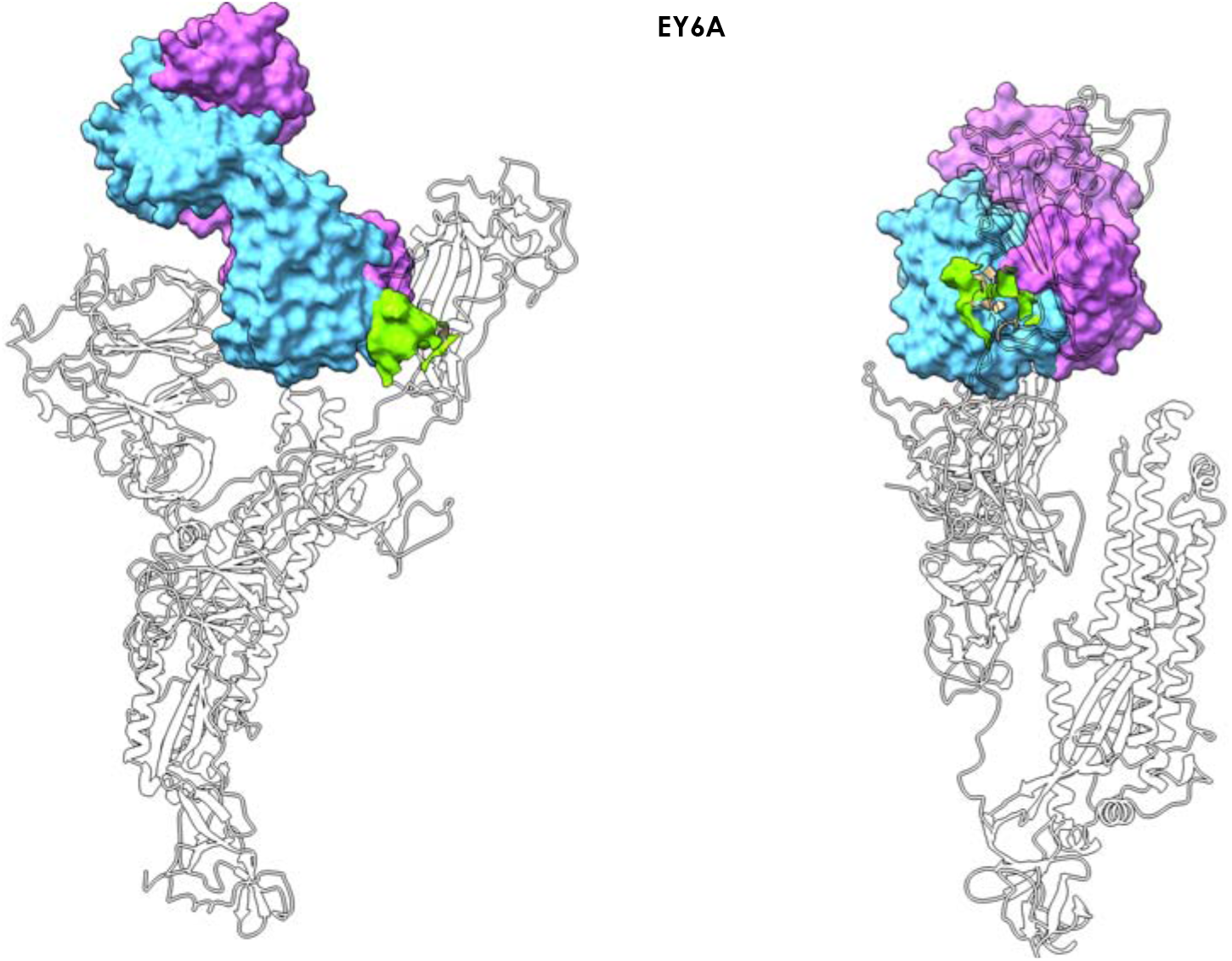
Antibody EY6A – Spike complex. The figure shows how Antibody **EY6A** (7byr.pdb) binds the RBD in the region of a cryptic epitope. The MLCE-predicted epitope region is shown in light green (lime) in two different orientations, indicating substantial contact formation with the antibody.

A second domain that is found to host a large network of non-optimized interactions corresponds to the N-terminal domain (Figure 1B, C, D). The latter has been shown to bind the new antibody **4A8** in a paper that was published during the preparation of this manuscript ^36^.

A third region predicted to be highly antigenic coincides with the central/C-terminal part of the S1A domain. In a recent cryo-EM study of polyclonal antibodies binding to the S protein, this substructure was shown to be in the vicinity of the density for **COV57** Fab(s), a novel Ab whose neutralizing activity showed no correlation with that of RBD-targeting Abs ^37^ (Figure 1B, D).

Furthermore, MLCE identifies a potentially highly reactive region in the S2 domain of the protein, in the CD region. This domain contains the epitope recently found to engage with 1A9 ^38^, an antibody recently shown to cross-react with S proteins of human, civet, and batcoronaviruses. This analysis also recognizes a potential antigenic region in a carbohydrate cluster located in the S2 domain of the protein: intriguingly, recent findings indicate that an oligosaccharide-containing epitope centered around this predicted region is targeted by the glycan-dependent antibody HIV-1 bnAb 2G12 ^39^ (Figure 1, 2).

The identification of energetically uncoupled domains also has mechanistic implications. Regions that are not involved in major intramolecular stabilizing interactions with the rest of the biomolecule can be displaced at minimal energetic costs, sustaining the large-scale conformational changes that typically underpin biological functionality. The boundary of the uncoupled N-terminal region (Figure 1, dark green domain) lies in physical proximity to the furin-targeted motif RRAR, essential for preactivation of SARS-CoV-2 Spike through proteolysis. In this framework, the uncoupling of a large part of the N-domain can synergize with (and favor through domain-displacement) the cleavage of specifically evolved furin motif to favor the detachment of S1-domain and the release of the S2 fusion machinery^8–10, 40^. Furthermore, the beta-sheet at the initial boundary of the C-terminal domain in the S2 (Red domain in Figure 1) is in close physical proximity to the fusion peptide (Figure 1D, E). It is tempting to suggest here that the release or conformational rearrangements of the C-terminal domain would be favored by its non-optimized interactions with the core of the stalk of the protein, optimally exposing the fusion peptide and favoring its integration with the host membrane ^40^.

Overall, these findings support the validity of our approach in identifying protein domains that can be aptly used as highly reactive immunogens, as they are most likely to be targeted by a humoral immune response. Our analysis predicts that regions other than the S protein RBD may represent alternative targets for neutralization or functional perturbation of SARS-CoV-2. On the one hand, this may be important considering the fact that RBD can also be the target on non-neutralizing antibodies, e.g. 30 2 2 ^35^. Indeed, using cocktails of antibodies to target different regions of S has recently been proposed as a viable therapeutic option ^36^.

Turning to our more stringent definition of epitope, based exclusively on the top 5% of the most weakly coupled residue pairs (5% cutoff), we next focus on those regions of the S antigen that can be involved in forming contacts with antibodies.

Importantly, one predicted conformational epitope with sequence (348)A-(352)A-(375)S-(434)IAWNS(438)-(442)DSKVGG(447)-(449)YNYL(452)-(459)S-(465)E-(491)PLQS(494)-(496)Q-(507)PYR(509) encompasses regions of the S protein in contact with antibodies **C105 (6xcn.pdb)**^37^, **S309**(6wpt.pdb; 6wps.pdb)^41^, **AB23** (7byr.pdb)^42^; **with nanobody H11-D4** (6z43.pdb); and with a recently reported synthetic nanobody (7c8v.pbd) (Figure 3).

Interestingly, an additional predicted patch comprising a set of decorating carbohydrates is correctly predicted to be part of the interface with antibody **S309 (6wpt.pdb; 6wps.pdb) ^41^,** with aminoacidic sequence (332)ITNLC(336)-(361)C and with (*N*334-linked) carbohydrate sequence β-D-GlcNAc–(α-L-Fuc)–β-D-GlcNAc–β-D-Man (i.e., UYB-(0fA)-4YB-VMB according to the *GLYCAM* naming convention ^43^). This predicted region sits notably close to the RBD interaction surface with ACE2.

Antibody **EY6A** (6zdh.pdb) binds the RBD in the region of the cryptic epitope described by Wilson and collaborators^35^ (Figure 4). Importantly, our predicted patch (365)YSVLYN(370)-(384)PTKLN(388) covers a significant part of the epitope. Once again, it is worth remarking that identification of this potentially immunoreactive patch is simply and exclusively obtained from structural and energetic interaction data generated for a protomer of the glycosylated, isolated S protein, after simulation with a general use forcefield (see Methods section).

With the more restrictive epitope prediction cutoff we clearly identify a reactive area in the N-terminal domain of the Spike protein. The predicted patch (184)GN(185)-(242)LAL(244)-(246)R-(248)Y-(258)WTAGA(262) contains residues R246 and W258 which were described as central determinants for contact between the N-terminal domain and antibody **4A8** ^36^ (Figure 5).

**Figure 5.**
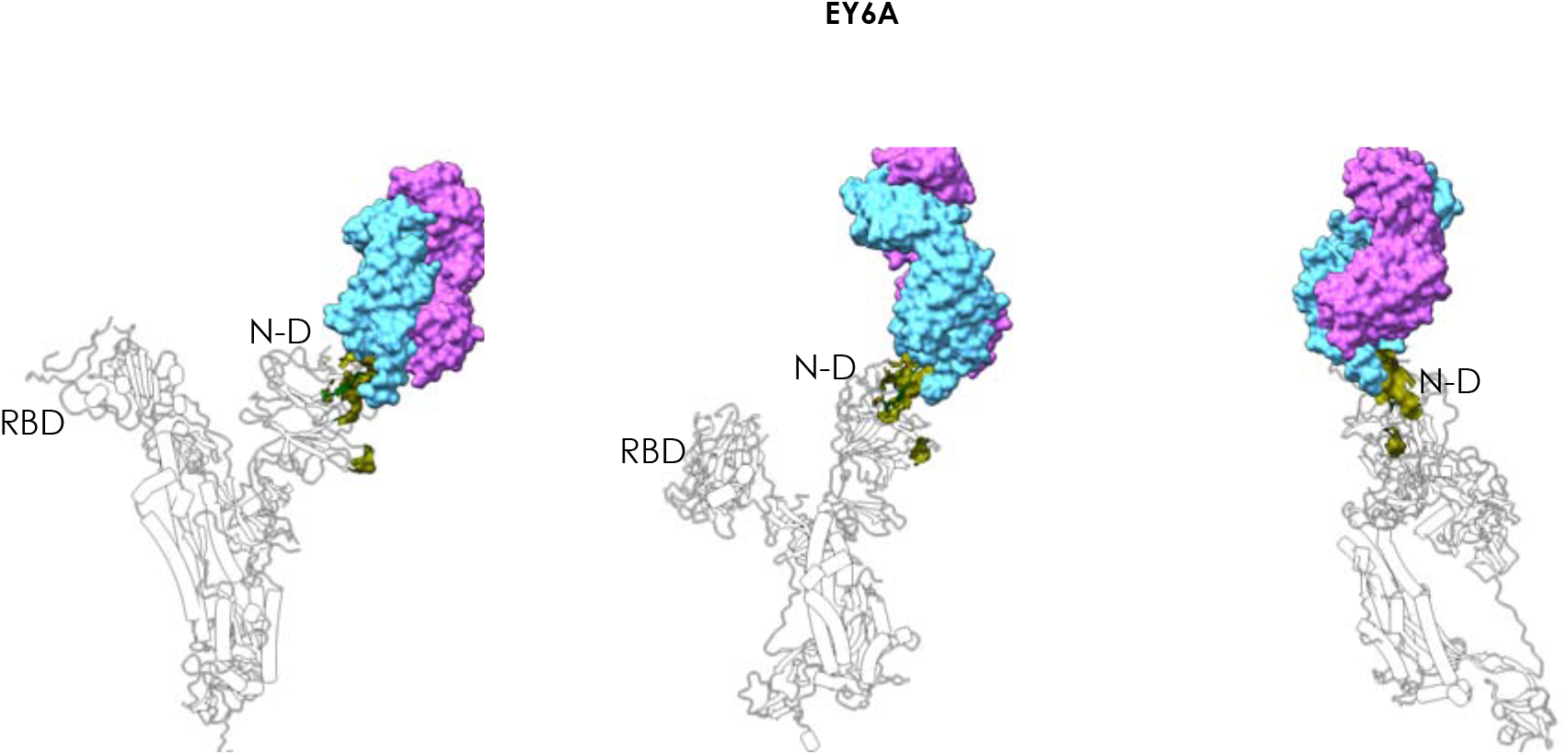
Antibody 4A8 – Spike complex. The figure shows how Antibody **4A8** binds the N-domain of Spike, supporting correct prediction of the epitope. The MLCE-predicted epitope region is shown in green in three different orientations, indicating substantial contact formation with the antibody. The Fab of the antibody is depicted as accessible surface in shades of blue.

Finally, the restrictive prediction identifies the sequence spanning residues 1076-1146, which includes amino acids 1111-1130, experimentally identified as the epitope for the monoclonal mAb **1A9** ^38^. Specifically our identified reactive sequence is the following: (1076) TTAPAICH(1083)-(1087)A-(1092)REG(1094)-(1096)FVSNGHWFVTQRN(1108)-(1112)P-(1114)I-(1116)T-(1118)DN(1119)-(1126)C-(1129)V-(1132)IVNNTVYDPLQELD(1146).

In general, our approach is able to identify potential immunoreactive domains and epitopes of the Spike protein based only on structural and energetic information. Sequences predicted to be reactive using the restrictive epitope definition (5% cutoff) can be used for generating optimized antigens in the form of synthetic peptide epitopes. Engineering such epitopes would entail the synthesis of conformationally preorganized peptidomimetics of the ‘natural’ reactive regions—with intra- and extracellular stability enhanced through, e.g, a combination of natural and non-natural aminoacids— which could reproduce the main structural and energetic conditions required to elicit a humoral immune response, as well as constituting candidates for vaccine development. Furthermore, reactive peptides thus identified may be suitable for use as baits in serologic diagnostic applications (e.g., in ELISA assays and in microarrays), to capture and detect to capture and detect not only circulating antibodies that are expressed in response to SARS-CoV-2 infections but also those that are endowed with neutralization activity and thus potentially predicting the infection outcome. As a further application, these peptide-based baits can represent a useful tool for isolating new mAbs and the screening of small molecules for drug development.

One the most significant aspects of our approach is that the S protein’s entire glycan shield is explicitly accounted for in the prediction of the immunoreactive regions. Indeed, the various oligosaccharide chains appear to behave differently (see differential coloring of oligosaccharide chains in Figure 1). In light of their stronger energetic coupling to other areas of the protein, some of the glycans are not recognized as epitopes, and thus form an integral part of the stabilizing intramolecular interaction network of S (white chains in Figure 1B); on the other hand, MLCE also identifies a second subset of poorly coupled oligosaccharides as potentially reactive epitopes (or part thereof) (colored oligosaccharide chains in Figure 1B; carbohydrate cluster in S2 targeted by the glycan-dependent antibody HIV-1 bnAb 2G12, see Figure 1B, 2), highlighting potential vulnerable spots in the glycan shield that could be exploited to design novel immunoreagents and vaccine candidates.

The portion of the glycan shield falling within the former category thus mainly serves to *protect* the protein from recognition by antibodies and consequently enhances viral infectiousness, as well as providing extra structural support. Two such glycans are further exemplified in **Figure 6**.

**Figure 6.**
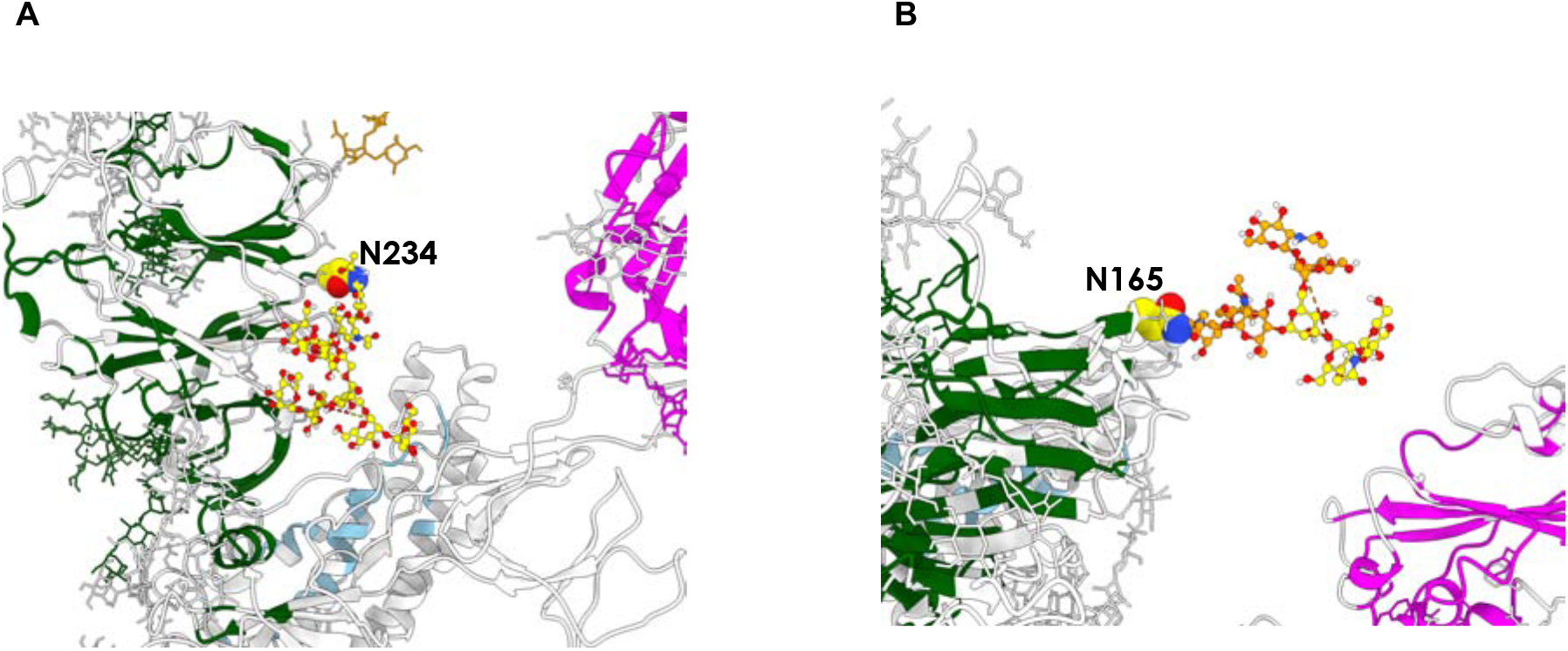
Glycans with different roles on Spike. **A)** The glycan chain attached to N234, which is predicted to be part of the networks of stabilizing interactions within the protein. **B)** The glycan chain attached to N165 is predicted to play a double role, a stabilizing one (yellow units) and an immunoreactive one (orange untis).

The first is the entire oligosaccharide fragment bound to N234 (Figure 6A), which is recognized by Amaro and coworkers as being crucial in ‘propping up’ the RBD^12^. Experimental deletion of *N*-glycans at this position by way of a mutation to Ala significantly modifies the conformational landscape of the protein’s RBD ^44^. The second is the portion of the N165-linked glycan whose subunits are rendered in yellow (**Figure 6B**): consistent with experimental studies indicating N165-linked oligosaccharides as structural modulators ^44^, we also find that the portion in question is *not* identified as a potential epitope, being consequently involved in diverting antibodies from targeting the region around N165 and thus preserving control of the S protein’s structural dynamics.

Reflecting the multifaceted roles of the glycan shield, the remaining part of the N165-linked glycan (**Figure 6B; orange**) appears instead to belong to the category of glycans that *are* potentially able to act as epitopes, since, unlike the part in yellow, we do detect it to be decoupled from the rest of the protomer.

It is particularly significant to underline that MLCE, whose physical basis is to identify non-optimized interaction networks, detects peptidic epitopes even when they are in proximity of (optimized, non-immunogenic) shielding carbohydrates. In light of this, it is reasonable to suggest that the protective effect of these particular carbohydrates may be circumvented and neutralized by exposing the underlying peptidic substructures. Furthermore, information on oligosaccharides identified as epitope constituents can be exploited to design glycomimics or glycosylated peptides as synthetic epitopes.

The latter aspect is indeed particularly relevant: small synthetic molecules that mimic antigenic determinants (and effectively act as their minimal surrogates) offer enticing opportunities to develop immunoreagents with superior characteristics in terms of ease of handling, reproducibility of batch-to-batch production, ease of purification, sustainable cost, and better stability under a variety of conditions. Furthermore, production of these molecules greatly reduces the risk of cross-reactivity with any copurified antigens, which is instead rife when dealing with recombinant proteins. In contrast to smaller peptides or sugar-decorated peptidomimetics, a full-length recombinant antigenic protein (or any protein-based detection device) would typically require more stringent conditions (e.g., in terms of temperature and humidity) for storage, transport, and management in order to preserve the protein in its properly folded active form. The same would be true for other vaccinal solutions such as deactivated pathogens.

Overall, our work confirms how simple and transparent structural and physico-chemical understanding of the molecule that is the key player in SARS-CoV-2 viral infection can be harnessed to guide the prediction of (in some cases experimentally confirmed) regions, that are involved in immune recognition and to understand its molecular bases. Agreement with experiment confirms that knowledge generated in the process has the potential of being translated into new molecules for vaccine and diagnostic development. In this context, we have also identified potentially reactive regions in the S protein stalk that are currently under experimental synthesis and testing.

Furthermore, the potential functional implications offered by the approach are illustrated by the fact that important domains/regions for the biological activation of the protein are naturally identified. This would give confidence for applying the method to understand the fine mechanistic determinants of possible new emerging mutants of the protein, expected to appear as the diffusion and host-adaptation of the virus progress. Finally, the possibility of accurately partitioning such a complex system in functional subunits could aptly be exploited in the parameterization of coarse-grained models for the description of the system at different length-scales.

This kind of structure-based computational approach can clearly expand the scope of simple structural analysis and molecular simulations. In applicative terms, generation of synthetic libraries based on predicted/identified epitopes (with possible addition of sugars) would definitely boost selection and screening of antigens for vaccine development.

## Methods

### STRUCTURE SELECTION FROM MOLECULAR DYNAMICS (RBD CLUSTERING)

Coordinates of the fully glycosylated SARS-CoV-2 S protein’s ‘RBD up’ protomer featured in this work originate from molecular dynamics (MD) simulations by Woods and coworkers ^13^, based on PDB ID: 6VSB. Throughout this work, we retain exactly the same forcefield parameters used by Woods *et al*. in their MD simulations: all residues except glycosylated asparagines are treated using the *ff14SB* forcefield, ^45^ whereas glycans and glycosylated asparagines are modeled using the *GLYCAM_06j* forcefield ^43^.

Clustering is based on root-mean-squared deviation of Cα atoms of the RBD domain in the ‘RBD up’ protomer, and performed with the *cpptraj* utility in *AmberTools* (version 17)^46^, after concatenating all six independent MD replicas ^13^ and aligning them with the ‘autoimage’ command. The chosen method is the Hierarchical Agglomerative Algorithm ^47^, with an epsilon value of 0.5. From each of the three most populated clusters, we isolate one representative frame, from which we retain the ‘RBD up’ protomer and its glycans, whilst again using *cpptraj* to discard all solvent molecules, ions, and the two ‘RBD down’ protomers. All subsequent calculations on these three ‘RBD up’ protomer models are listed chronologically in the subsections that follow.

### MINIMZATION

A 200-step minimization of each of the three protomer models is carried out using the default procedure (i.e., steepest descent for 10 steps; then conjugate gradient) implemented in the MD engine *sander* in the *AMBER* software package (version 18))^46^. Protomers are minimized using the generalized Born (GB) implicit solvent model as parametrized by Onufriev et al. ^48^, with a universal 12.0 Å cutoff applied in the calculation of Lennard-Jones and Coulomb interactions (neither of which are calculated beyond this limit). For this stage, concentration of (implicit) mobile counterions in the GB model is set to 0.1 M, and the solvent-accessible surface area (SASA) is computed according to the LCPO method (linear combinations of pairwise overlaps) ^49^.

### MM/GBSA CALCULATIONS

MM/GBSA calculations ^33^ are performed on each of the three minimized ‘RBD up’ protomers using the dedicated mm_pbsa.pl utility in *AmberTools* (version 17). The purpose of these calculations is to obtain a breakdown of nonbonded energy interactions (i.e., electrostatic, van der Waals, implicit solvation contributions and, in this case, 1-4 interactions) between every possible pair of residues in the protomer (aminoacids and monosaccharides alike): for a protomer composed of *N* residues, this leads to a symmetric ***N × N*** interaction matrix ***M_ij_***.^50–51^

The implicit GB solvation model used in these calculations is identical to the one used in the preceding minimization step (vide supra), except that the implicit ion concentration is set to 0.0 M, and SASA is computed with the ICOSA method (based on icosahedra around each atom that are progressively refined to spheres).

### ENERGY DECOMPOSITION

The symmetric interaction matrix ***M_ij_*** obtained from separate MM/GBSA calculations (vide supra) on each of the three S protein protomer models under study (with ***N*** residues) can be expressed in terms of its eigenvalues and eigenvectors as

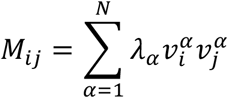

where ***λ_α_*** is the a-th eigenvalue and 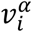 is the *i*^th^ component of the corresponding eigenvector.

It was previously shown in a number of cases that **eigenvector** (***v*^1^**), also called *first eigenvector*, associated with the lowest eigenvalue *λ*_1_. allows to identify most of the crucial aminoacids necessary for the stabilization of a protein fold, and consequently those aminoacids that are minimally coupled to such core. The latter were shown to correspond to potential interaction regions.

In the case of multidomain proteins such as S, the first eigenvector is not sufficient, and more eigenvectors are needed to capture the essential interactions for folding/stability and binding. The interaction matrix ***M_ij_*** is thus decomposed instead via the alternative approach developed by Genoni *et al*.^34^. In this scenario, the aim is to select the smallest set of ***N_e_*** eigenvectors that cover the largest part of residues (i.e., components) with the minimum redundancy under the assumption that: (a) for each domain there should exist only one associated eigenvector recapitulating its most significant interactions; (b) each “domain eigenvector” has a block structure whereby its significant components correspond to the residues belonging to the identified domain; (c) combination of all significant blocks covers all residues in the protein. Matrix ***M_ij_*** can thus be reformulated as a simplified matrix 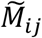:

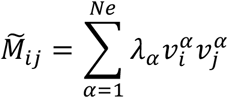

(where this time the sum occurs over ***N_e_*** essential eigenvectors instead of ***N*** residues). As detailed by Genoni *et al*. ^34^., the essential folding matrix 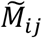 is subsequently further filtered through a symbolization process to emphasize the significant non-bonded interaction, yielding 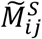 and finally subjected to a proper clustering procedure leading to domain identification.

The final simplified matrix 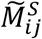 resulting from domain decomposition thus only reports those residue pairs in the protomer that exhibit the strongest and weakest energetic interactions.

### MLCE

Final epitope predictions are made using the Matrix of Local Coupling Energies method (MLCE), in which analysis of a given protein’s energetic properties is combined with that of its structural determinants ^21, 23^. This approach allows to identify nonoptimized, contiguous regions on the protein surface that are deemed to have minimal coupling energies with the rest of the structure, and that have a greater propensity for recognition by Abs or other binding partners.

The MLCE procedure entails cross-comparison of the simplified pairwise residue-residue energy interaction matrix 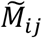 resulting from domain decomposition (vide supra) with a pairwise residueresidue contact matrix ***C_ij_***. The latter matrix namely considers a pair of residues to be spatially contiguous (i.e., ‘in contact’) if they are closer than an arbitrary 6.0 Å-threshold; contact distances are measured between Cβ atoms in the case of non-glycine aminoacid residues, H atoms in the case of glycine residues, and between C1 atoms in the case of glycan residues.

The Hadamard product of the two matrices yields the matrix of the local pairwise coupling energies ***MLCE_ij_***:

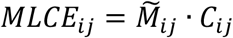

Deriving the MLCE matrix allows to rank spatially contiguous residue pairs with respect to the strength of their energetic interactions (weakest to strongest). Selection of proximal pairs showing the weakest coupling with the rest of the protein ultimately defines putative epitopes; two distinct selections are carried out on the basis of two possible weakness (softness) cutoffs (5% or 15%), corresponding to the top 5% or 15% spatially contiguous residue pairs with the lowest energetic interactions.

## Acknowledgement

The authors thank Prof. Robert J. Woods and Dr. Oliver Grant (University of Georgia) for kindly providing atomistic molecular dynamics simulations of the fully glycosylated spike proteins. This research was supported by a Grant from “Cassa di Risparmio di Padova e Rovigo (CARIPARO) PROGETTI DI RICERCA SUL COVID-19” to GC and AR.

## Notes

### Competing Interest Statement

The authors have declared no competing interest.

